# Comparative Analysis of *UL16* Mutants Derived from Multiple Strains of HSV-2 and HSV-1 Reveals Species-Specific Requirements for the UL16 Protein

**DOI:** 10.1101/274548

**Authors:** Jie Gao, Xiaohu Yan, Bruce W. Banfield

## Abstract

Orthologs of the herpes simplex virus (HSV) *UL16* gene are conserved throughout the *Herpesviridae*. Because of this conservation, one might expect that these proteins perform similar functions for all herpesviruses. Previous studies on a *UL16* null mutant derived from HSV-2 strain 186 revealed a roughly 100-fold replication defect and a critical role for UL16 in the nuclear egress of capsids. These findings were in stark contrast to what has been observed with *UL16* mutants of HSV-1 and pseudorabies virus where roughly 10-fold replication deficiencies were reported that were accompanied by defects in the secondary envelopment of cytoplasmic capsids. One possible explanation for this discrepancy is that the HSV-2 186 strain is not representative of the HSV-2 species. To address this possibility, multiple *UL16* null mutants were constructed in multiple HSV-2 and HSV-1 strains by CRISPR/Cas9 mutagenesis and their phenotypes characterized side-by-side. This analysis showed that all the HSV-2 *UL16* mutants had 50 to 100-fold replication deficiencies that were accompanied by defects in the nuclear egress of capsids as well as defects in the secondary envelopment of cytoplasmic capsids. By contrast, most HSV-1 *UL16* mutants had 10-fold replication deficiencies that were accompanied by defects in secondary envelopment of cytoplasmic capsids. These findings indicated that UL16 has HSV species-specific functions. Interestingly, HSV-1 UL16 could promote the nuclear egress of HSV-2 *UL16* null strains, suggesting that, unlike HSV-1, HSV-2 lacks an activity that can compensate for nuclear egress in the absence of UL16.

**Importance:** HSV-2 and HSV-1 are important human pathogens that cause distinct diseases in their hosts. A complete understanding of the morphogenesis of these viruses is expected to reveal vulnerabilities that can be exploited in the treatment of HSV disease. UL16 is a virion structural component that is conserved throughout the *Herpesviridae* and functions in virus morphogenesis, however, previous studies have suggested different roles for UL16 in the morphogenesis of HSV-2 and HSV-1. This study sought to resolve this apparent discrepancy by analyzing multiple UL16 mutant viruses derived from multiple strains of HSV-2 and HSV-1. The data indicate that UL16 has HSV species-specific functions insofar as HSV-2 has a requirement for UL16 in the escape of capsids from the nucleus whereas both HSV-2 and HSV-1 require UL16 for final envelopment of capsids at cytoplasmic membranes.

## Introduction

While the early stages of herpesvirus assembly take place in the nucleus, the final stages of virion assembly occur in the cytoplasm of infected cells. Viral DNA is packaged into preformed procapsids in the infected cell nucleus resulting the formation of C-capsids that are competent for subsequent stages in virion maturation. To reach the cytoplasm, genome containing C-capsids must transit across the inner and outer nuclear membranes utilizing a process referred to as nuclear egress - a subject of recent and intense investigation by numerous laboratories (1, 2). Nuclear egress of C-capsids occurs through primary envelopment of capsids at the inner nuclear membrane followed by de-envelopment, and release of capsids into the cytoplasm, through fusion of the perinuclear virion envelope with the outer nuclear membrane. Once in the cytoplasm, the C-capsid acquires its final envelope by budding into membrane vesicles derived from the trans-Golgi network, or an endocytic compartment, in a process referred to as secondary envelopment (3-5). Finally, enveloped virions contained within vesicles are transported to the cell surface where they fuse with the plasma membrane releasing the mature virion into the extracellular space.

This study concerns the functions of the HSV UL16 protein in virion assembly. Orthologs of the HSV UL16 protein are conserved throughout the *Herpesviridae*, however its specific roles in the virus replicative cycle are poorly understood. Contributing to this lack of clarity are seemingly conflicting reports on the functions of the HSV-2 and HSV-1 UL16 orthologs. This laboratory recently reported that deletion of *UL16* from the HSV-2 186 strain resulted in a roughly 100-fold reduction in virus replication and a failure of C-capsids to undergo efficient nuclear egress (6). In contrast to these findings, several groups have reported that HSV-1 *UL16* mutants have more modest, roughly 10-fold, replication deficiencies and are defective in secondary envelopment rather than nuclear egress (7, 8). It is noteworthy that studies on the UL16 ortholog from pseudorabies virus (PRV), a virus distantly related to HSV, closely resembled the findings seen with HSV-1, where roughly 10-fold replication deficiencies associated with defective secondary envelopment were reported (9). What could explain these conflicting reports? One possibility was that the single strain of HSV-2 studied by Gao and colleagues (6), strain 186, was an outlier and the results obtained with this strain were not representative of the HSV-2 species as a whole. Another possibility was that the HSV-1 and PRV strains analyzed previously were constructed in such a way that promoted the selection of suppressor mutations that might overcome the replication deficiencies and nuclear egress phenotypes exhibited by HSV-2 (186) Δ16, which, by contrast, was isolated on complementing cells to mitigate the selection of suppressor mutations. A third possibility was that UL16 has species-specific functions during the replication of HSV-2 and HSV-1.

The goal of this study was to resolve this controversy by performing a side-by-side analysis of a panel of newly constructed *UL16* mutants derived from multiple strains of HSV-2 and HSV-1. All strains were constructed using the same procedures. CRISPR/Cas9 based mutagenesis was used to create the mutant virus genomes and all *UL16* mutant viruses were propagated on UL16-expressing cells to avoid the enrichment of suppressor mutants during strain isolation. To extend the previous analysis of HSV-2 strain 186 we chose build *UL16* mutants into strains HG52 and SD90e. Strain HG52 is a well-studied HSV-2 reference strain that was the first to be completely sequenced (10), whereas strain SD90e is a low passage clinical isolate that has been proposed to serve as a new HSV-2 reference strain (11, 12). For the construction of new HSV-1 *UL16* mutants, we chose to utilize strains F and KOS, two well-studied laboratory strains that have been used by others to study the function of UL16 (7, 8). Our analysis indicates that UL16 plays a critical role in both the nuclear egress and secondary envelopment of HSV-2 strains whereas HSV-1 UL16 functions primarily in the secondary envelopment of cytoplasmic capsids. Interestingly, trans-complementation experiments revealed that HSV-2 and HSV-1 UL16 can substitute for each other suggesting that the genetic basis for the species-specific requirements of *UL16* reside outside the *UL16* locus.

## Results

### Construction of HSV *UL16* mutant viruses

We constructed *UL16* deletions in multiple HSV-2 and HSV-1 strains to enable a comparative analysis of these viruses. To avoid any selective pressure during the isolation of these strains UL16 expressing cell lines were used. *UL16* deletions were constructed in HSV-2 strains SD90e and HG52, and HSV-1 strains KOS and F by CRISPR/Cas9-based mutagenesis as described in Materials and Methods. Two mutants derived independently from each strain were selected for further study and all mutants studied were used at low passage. DNA sequencing of HSV-2 and HSV-1 deletion mutants revealed the exact nature of the *UL16* mutants isolated (Fig. 1A and 1B). To verify that the *UL16* mutants did not express UL16, Vero cells were infected with the different HSV-2 and HSV-1 *UL16* mutants at MOI of 0.5 and cells lysates prepared at 24 hpi. Western blots of cell lysates were probed for HSV-2 and HSV-1 UL16, the immediate early protein ICP27 (infection control) and β-actin (loading control) (Fig. 1C and 1D). Full-length UL16 was observed in all wild type virus infected cell lysates, while no full-length UL16 protein was expressed in lysates from any *UL16* mutant infected cells. Notably, UL16SΔ10-59 and UL16KΔ6-139 infected cell lysates, contained truncated UL16 proteins identified by asterisks in Figs.1C and D. These data confirmed that all *UL16* mutants failed to produce full length UL16 protein.

**Fig 1.**
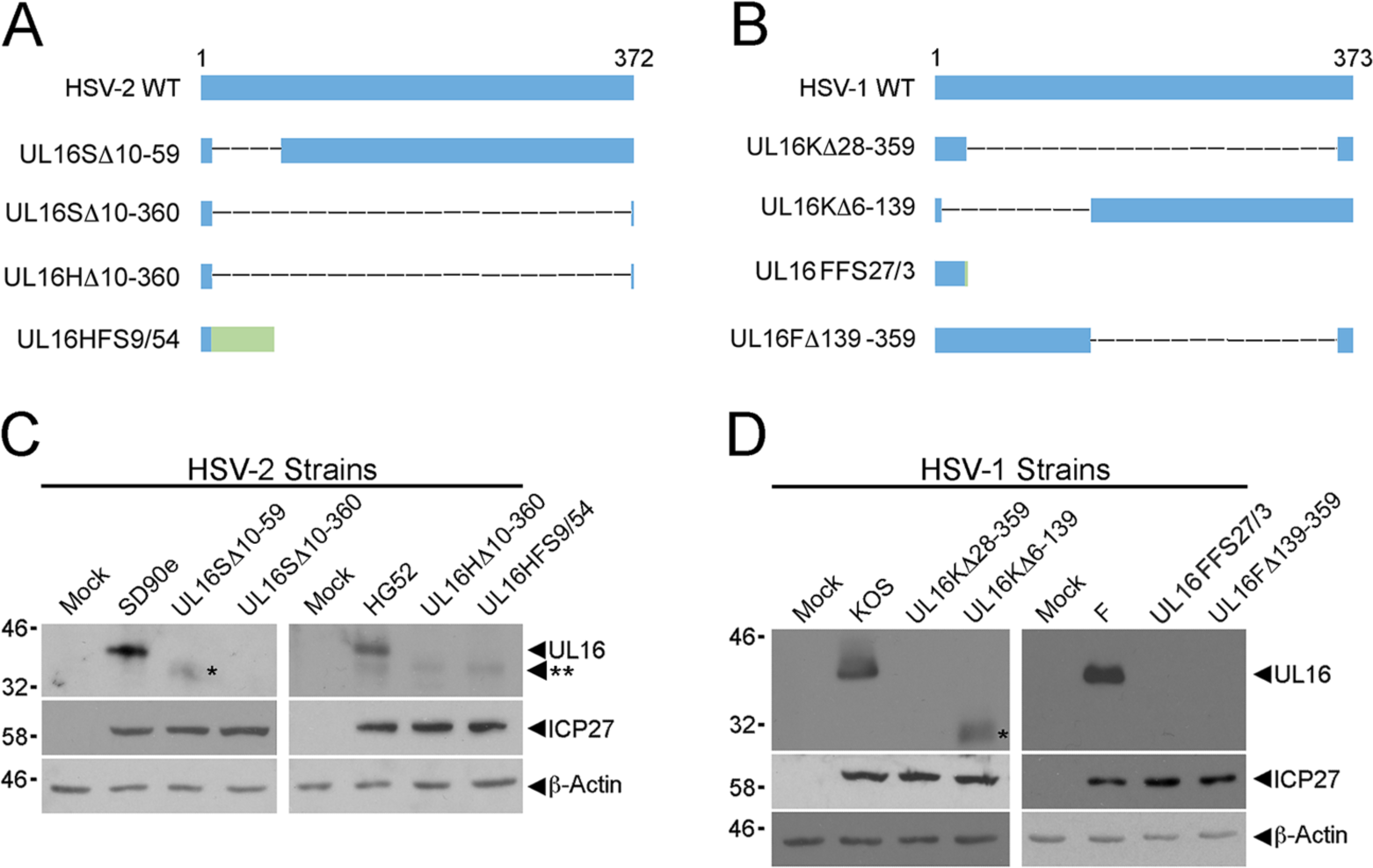
HSV *UL16* mutants. Diagrams of full-length (WT) and mutant UL16 proteins from HSV-2 (**A**) and HSV-1 (**B**) are shown. Four *UL16* deletion mutants of each HSV species were selected for further analysis. Blue bars indicate UL16 protein sequence, dashed lines represent deleted sequences and green bars indicate non-UL16 amino acids that arise due to frame shift. In the nomenclature used, the first letter after UL16 indicates the parental strain (S= SD90e; H= HG52; K= KOS, F= F); Δ refers to an in-frame deletion and the numbers following Δ indicate the position of the codons that were deleted from the *UL16* gene; FS refers to a frame shift and the first number following FS refers to the position of the codon where the frame shift occurred and the last number refers to the number of non-UL16 amino acids. Western blots of cell lysates from Vero cells infected with the strains shown in **A** and **B** were probed using antiserum against HSV -2 UL16 (**C**) or HSV-1 UL16 (**D**). ICP27 antiserum was used as a positive control for viral infection, while β-actin was used as a loading control. Single asterisks indicate truncated forms of UL16. Double asterisk indicates position of a non-specific band detected in HG52 infected cell lysates in panel C.

### UL16 is required for efficient cell-to-cell spread of both HSV-2 and HSV-1 strains

To determine if the cell-to-cell spread properties of these new *UL16* deletion mutants were consistent with our results with HSV-2 strain 186 (6) and those reported by others for HSV-1 strains KOS and F (8), monolayers of L, L16 (express HSV-2 UL16), L16K (express HSV-1 UL16) or Vero cells were infected with *UL16* mutants and their parental viruses (Fig. 2). At 72 hpi, cells were fixed and stained with methylene blue. All HSV-2 *UL16* null mutants formed visible plaques on complementing L16 cells, but did not form visible plaques on non-complementing L cells (Fig. 2A). Similarly, all HSV-1 *UL16* mutants formed visible plaques on complementing L16K cells, but not on L cells (Fig. 2C). Importantly, all *UL16* null strains formed visible plaques on non-complementing Vero cells, albeit much smaller than those formed in comparison to their parental strains (Fig. 2A and 2C), indicating some capacity for spread between Vero cells that was not seen on L cell monolayers. These data suggested that HSV-1 and HSV-2 *UL16* mutants have similar deficiencies in cell-to-cell spread.

**Fig 2.**
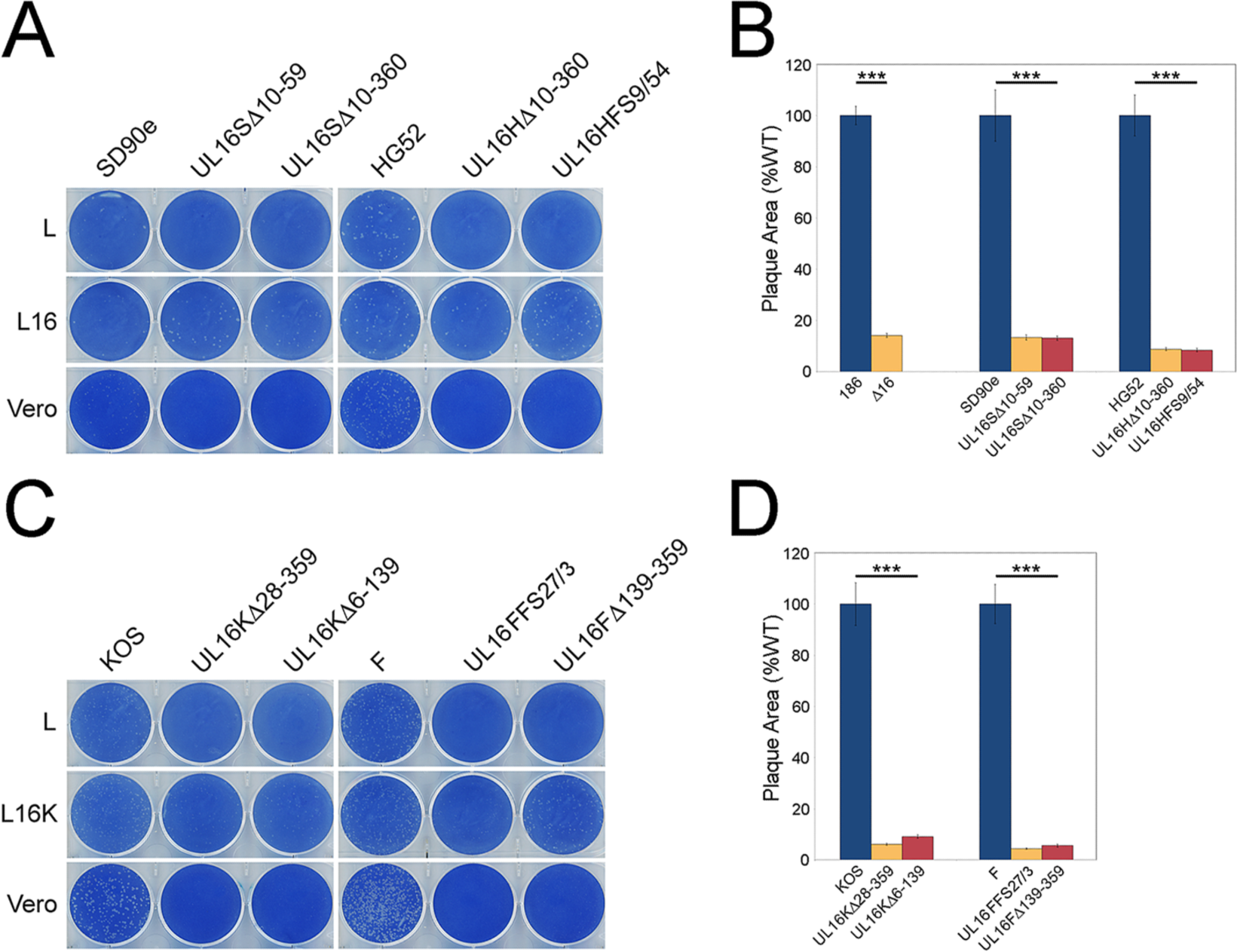
Cell-to-cell spread capabilities of HSV *UL16* mutants. **(A)** Identical dilutions of each HSV strain were used to infect the non-complementing and complementing cell monolayers indicated. Cells were fixed and stained with 0.5% methylene blue in 70% methanol at 72 hpi. **(B)** Vero cells were infected with the indicated viruses, and cells were fixed and plaques stained using antiserum against HSV Us3 by indirect immunofluorescence microscopy at 24 hpi. Plaque sizes were determined as described in Materials and Methods (n=40 plaques per strain). Error bars represent standard error of means. HSV wild type strains 186, SD90e, HG52, KOS and F were normalized to 100%. (* * *, *P*<0.0001).

To quantify the abilities of *UL16* deletion viruses to spread, we measured the area of the plaques produced by the *UL16* deletion mutants on non-complementing Vero cells. At 24 hpi plaques were fixed and stained using antisera against HSV Us3 and the areas of the plaques measured using ImagePro 6.3. The two HSV-2 (SD90e) *UL16* mutants formed plaques approximately 13% of the size of their parental strain, while HSV-2 (HG52) *UL16* mutant plaques were around 8% the size of WT HG52 plaques (Fig. 2B). In addition, the plaque size of our original HSV-2 186 strain *UL16* null mutant, Δ16, was 14% that of WT 186 strain (Fig. 2B). Surprisingly, all HSV-1 *UL16* null mutants formed plaques roughly 95% smaller than their parental strains, similar to what was observed with HSV-2 *UL16* null strains (Fig. 2D). Collectively, these findings suggested that UL16 is critical for virus spread on non-complementing cells and no obvious differences between HSV-2 and HSV-1 virus spread were observed in the absence of UL16.

### Replication kinetics of HSV-2 and HSV-1 *UL16* null strains

To provide a more comprehensive view of the replication defects of HSV-2 and HSV-1 *UL16* deletion mutants, we performed multi-step growth analysis. Monolayers of Vero cells were infected with HSV-2 and HSV-1 *UL16* mutants and their corresponding parental strains at an MOI of 0.01. Cells and medium were harvested together at indicated time points after infection and titrated on monolayers of complementing L16 cells. The results showed that the HSV-2 (SD90e and HG52) *UL16* deletions had approximately 100-fold and 50-fold reductions in end-point titres compared their parental strains, respectively (Fig. 3). By contrast, with one exception, our KOS and F *UL16* mutants had roughly 10-fold reductions in virus replication compared to their parental strain (Fig. 3). UL16FFS27/3 was an outlier insofar as it replicated much more poorly (400-fold lower than WT F) than the other HSV-1 strains analyzed. With the exception of the UL16FFS27/3 strain, these data are consistent with previous findings (6-8) indicating that HSV-2 *UL16* mutants replicate less efficiently than HSV-1 *UL16* mutants.

**Fig 3.**
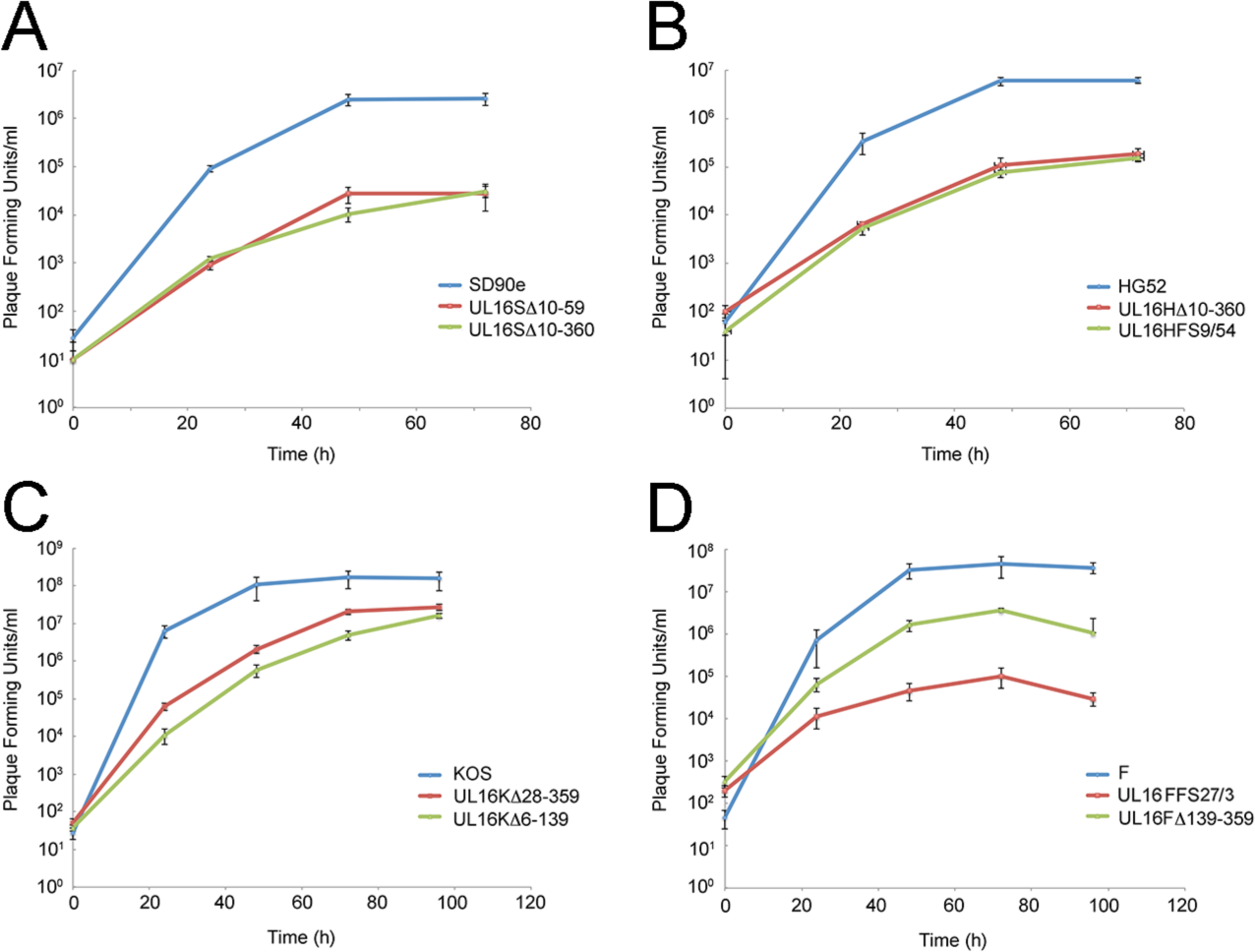
Replication kinetics of HSV *UL16* deletion mutants. Monolayers of Vero cells were infected with parental HSV strains **(A.** SD90e, **B.** HG52**, C.** KOS, and **D.** F) and their corresponding *UL16* deletion mutants at an MOI of 0.01. Cells and medium were harvested together at indicated times post infection, and titrated on monolayers of L16 cells. Each data point represents the average data from two biological replicates, each of which was titrated in triplicate. Error bars are standard errors of the means.

### Reciprocal complementation between HSV-2 and HSV-1 UL16

To examine whether HSV-2 and HSV-1 UL16 proteins could functionally compensate for each other, reciprocal complementation assays were performed. Monolayers of Vero, Vero16 (expressing HSV-2 UL16) and Vero16K (expressing HSV-1 UL16) were infected with the same dilutions of HSV-2 and HSV-1 *UL16* deletion viruses and their parental strains. At 72 hpi cells were fixed and stained with methylene blue. Interestingly, all HSV-2 and HSV-1 *UL16* mutants formed plaques on Vero, Vero16 and Vero16K (Fig. 4). Notably, *UL16* mutants formed very small plaques on Vero cells compared to their parental strains, consistent with the data shown above (Fig. 2). The data indicate that the HSV-2 UL16 protein can complement HSV-1 *UL16* null strains (Fig. 4A) and vice versa (Fig. 4B).

**Fig 4.**
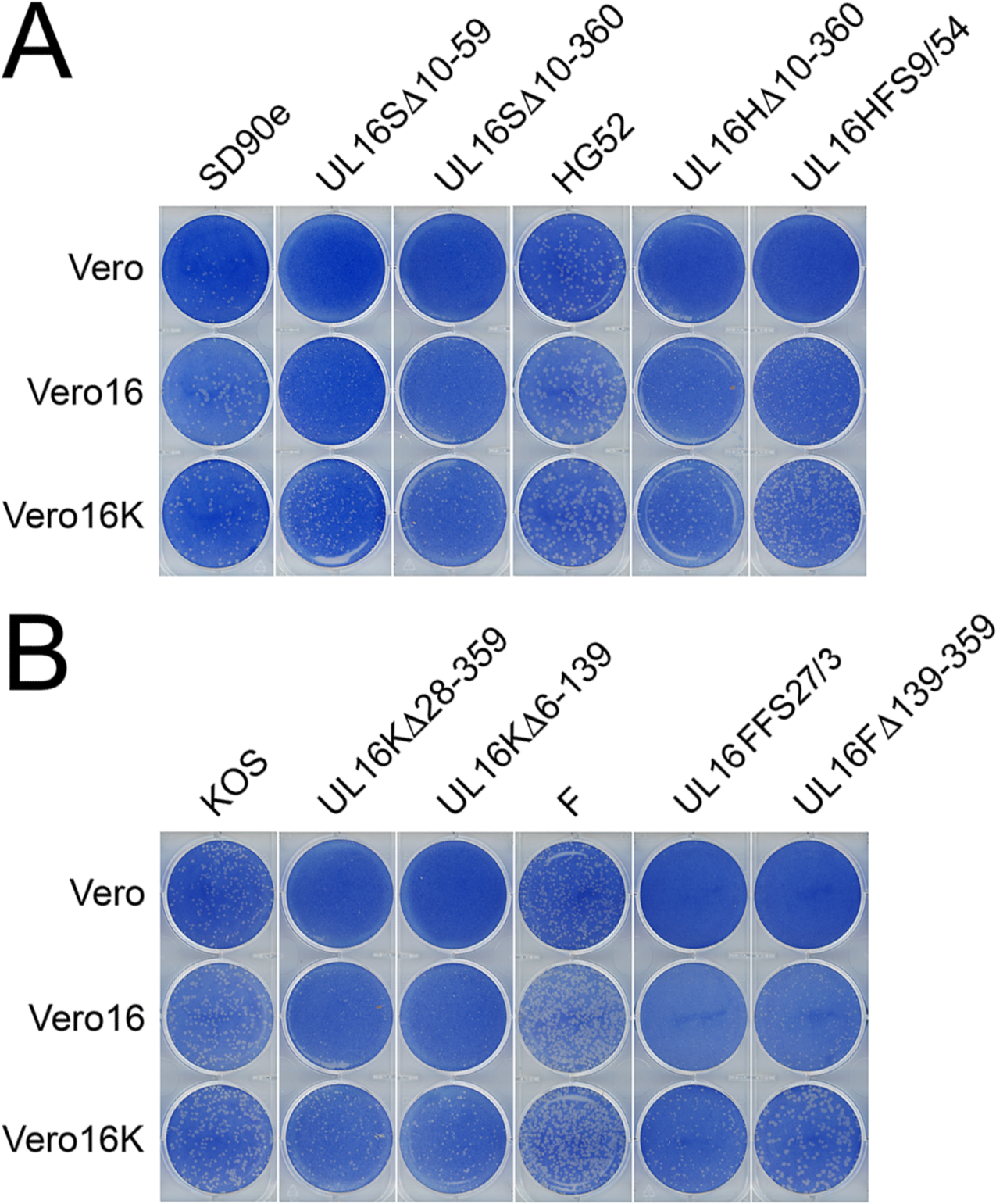
Reciprocal complementation between HSV-2 and HSV-1 UL16. Monolayers of Vero, Vero16 and Vero16K cells were infected with identical dilutions of each HSV-2 (**A**) and HSV-1 (**B**) strain. At 72 hpi cells were fixed and stained with 0.5% methylene blue in 70% methanol.

### Species specific requirements for the UL16 protein

To define and compare the stages at which our HSV-2 and HSV-1 *UL16* mutants were blocked in their maturation, transmission electron microscopy (TEM) was performed. Vero cells were infected with HSV-1 and HSV-2 *UL16* mutants and corresponding parental strains, and cells were fixed and processed for TEM at 16 hpi as described in Materials and Methods. A, B and C-capsids were readily observed in the nuclei of parental HSV-2 (SD90e) infected cells (Fig. 5A), and UL16SΔ10-360 infected cells (Fig.5C). However, similar to what we reported previously for HSV-2 (186) (6), many more cytoplasmic capsids were observed in HSV-2 (SD90e) infected cells (Fig.5B) than in cells infected with its corresponding *UL16* mutant, UL16SΔ10-360 (Fig.5D). Numerous capsids were observed both in the nuclei and cytoplasm of HSV-1 (F) and UL16FΔ139-359 infected cells (Fig. 6). However, fewer enveloped cytoplasmic capsids were observed in cells infected with the *UL16* mutant, UL16FΔ139-359 (Fig. 6D), compared to HSV-1 (F) infected cells (Fig. 6B). These findings are consistent with previous reports indicating that HSV-1 UL16 functions in secondary envelopment (8).

**Fig 5.**
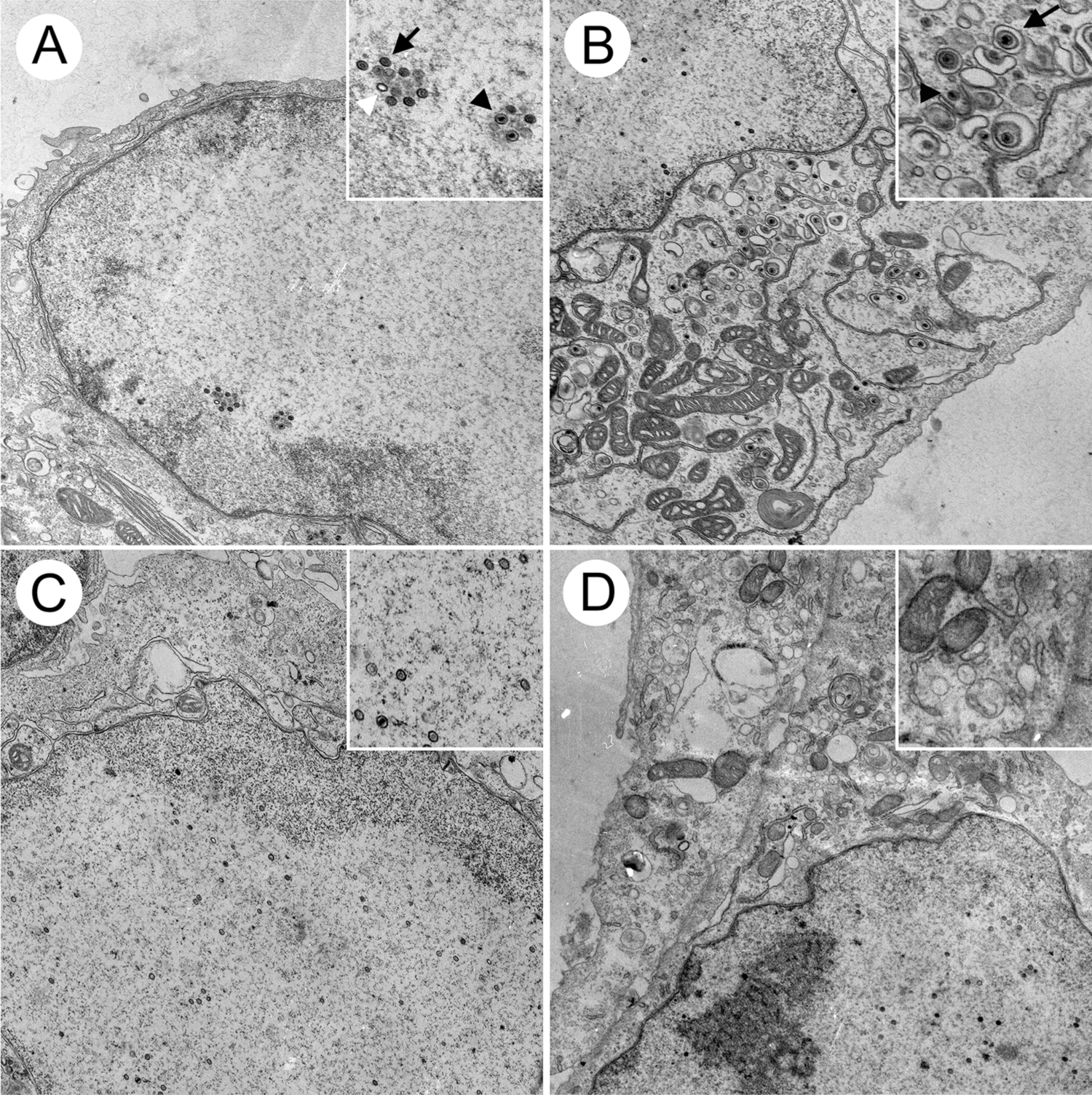
Ultrastructural analysis of HSV-2 infected cells. Vero cells were infected with HSV-2 SD90e (**A** and **B**) and the *UL16* deletion mutant, UL16SΔ10-360 (**C** and **D**), at an MOI of 3. At 16 hpi, cells were fixed and processed for TEM as described in Materials and Methods. Non-enveloped cytoplasmic capsids and enveloped virions can be observed in the cytoplasm of SD90e infected Vero cells (**B**). These structures were rarely observed in the cytoplasm of UL16SΔ10-360 infected cells (**D**). Nuclear capsids were readily detected in the nuclei of SD90e (**A**) and UL16SΔ10-360 (**C**) infected cells. Insets in each panel show magnified portions of the images. White arrowhead in the panel A inset identifies an A capsid, whereas a B capsid is identified with a black arrow and a C capsid identified with a black arrowhead. Inset to panel B identifies an enveloped capsid with a black arrow and a non-enveloped capsid with a black arrowhead.

**Fig 6.**
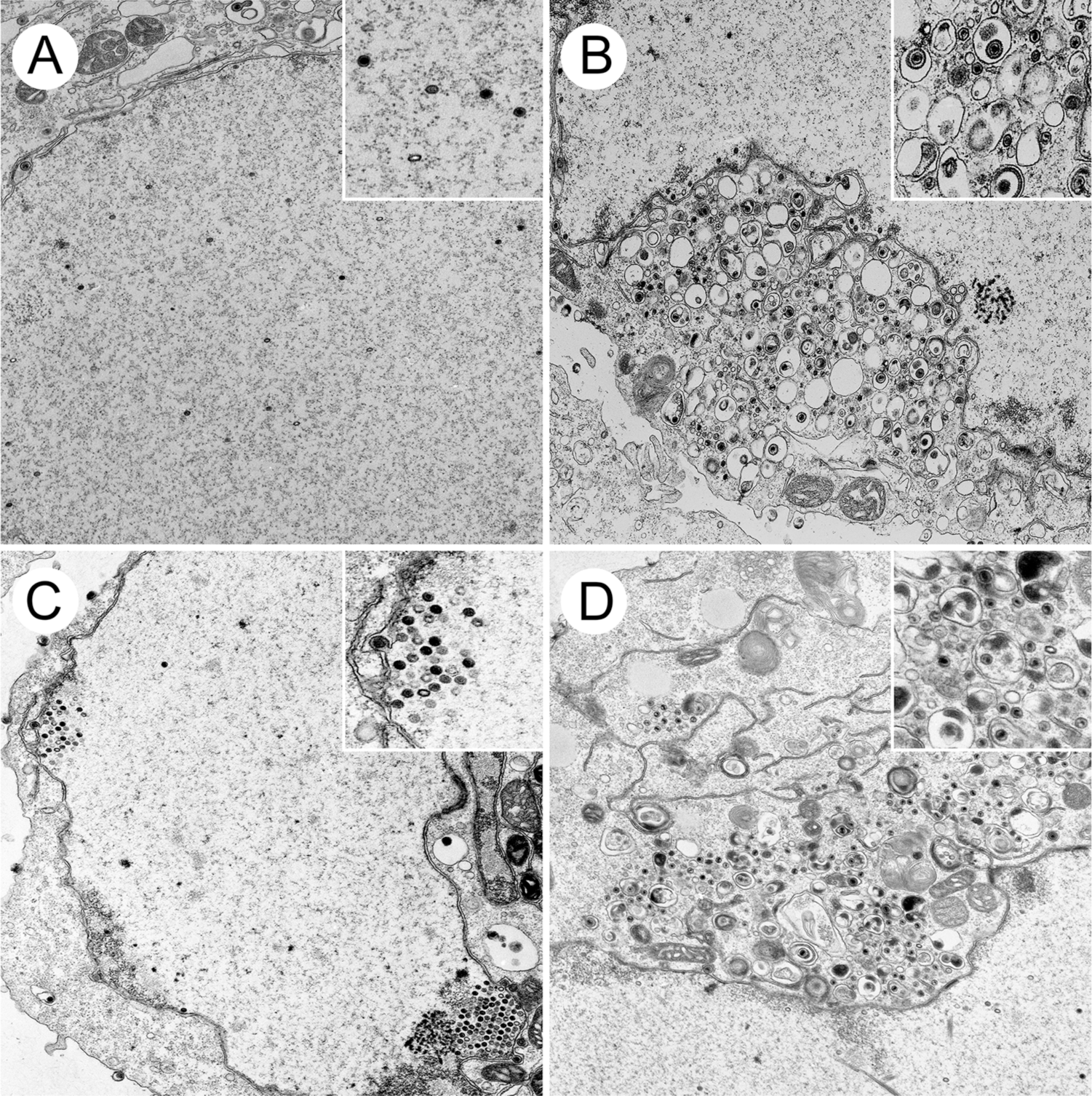
Ultrastructural analysis of HSV-1 infected cells. Vero cells were infected with HSV-1 F (**A** and **B**) and the *UL16* deletion mutant, UL16FΔ139-359 (**C** and **D**), at an MOI of 3. At 16 hpi, cells were fixed and processed for TEM as described in Materials and Methods. Non-enveloped cytoplasmic capsids and enveloped virions can be observed in the cytoplasm of F infected cells (**B**). Enveloped virions were less frequently observed in the cytoplasm of UL16FΔ139-359 infected cells where non-enveloped capsids were abundant (**D**). Nuclear capsids were readily detected in the nuclei of both F (**A**) and UL16FΔ139-359 (**C**) infected cells.

To quantify the distribution of capsids in the presence and absence of UL16, viral particles at various stages of maturation were classified and counted in ten independent images of Vero cells infected with the strains listed in Table 2 and the ratios of intranuclear C-capsids:cytoplasmic capsids and enveloped:cytoplasmic capsids were analyzed in more detail (Fig. 7). We chose to focus on intranuclear C-capsids, instead of A, B and C-capsids together, because C-capsids are preferentially selected for primary envelopment(13). The ratio of intranuclear C-capsids:cytoplasmic capsids was significantly greater in cells infected with HSV-2 *UL16* mutants than in cells infected with their parental counterparts (Fig. 7A). By contrast, the ratio of intranuclear C-capsids:cytoplasmic capsids was greater for parental HSV-1 strains than for their *UL16* mutants suggesting that HSV-1 UL16 does not play a discernable role in nuclear egress. The fact that the ratios of intranuclear C-capsids:cytoplasmic capsids were significantly lower for HSV-1 *UL16* mutants compared to their parental strains may be due to the accumulation of cytoplasmic capsids in cells infected with the *UL16* mutant strains (Table 2 and Fig.6D). The mean ratios of enveloped/cytoplasmic capsids for HSV-2 and HSV-1 *UL16* mutant strains were significantly lower than their parental strains indicating that UL16 functions in secondary envelopment for both species of HSV (Fig. 7B). Taken together these data indicate that UL16 has species-specific functions in HSV infection such that HSV-2 relies strongly on UL16 for nuclear egress whereas both HSV-2 and HSV-1 rely on UL16 for efficient secondary envelopment.

**Fig 7.**
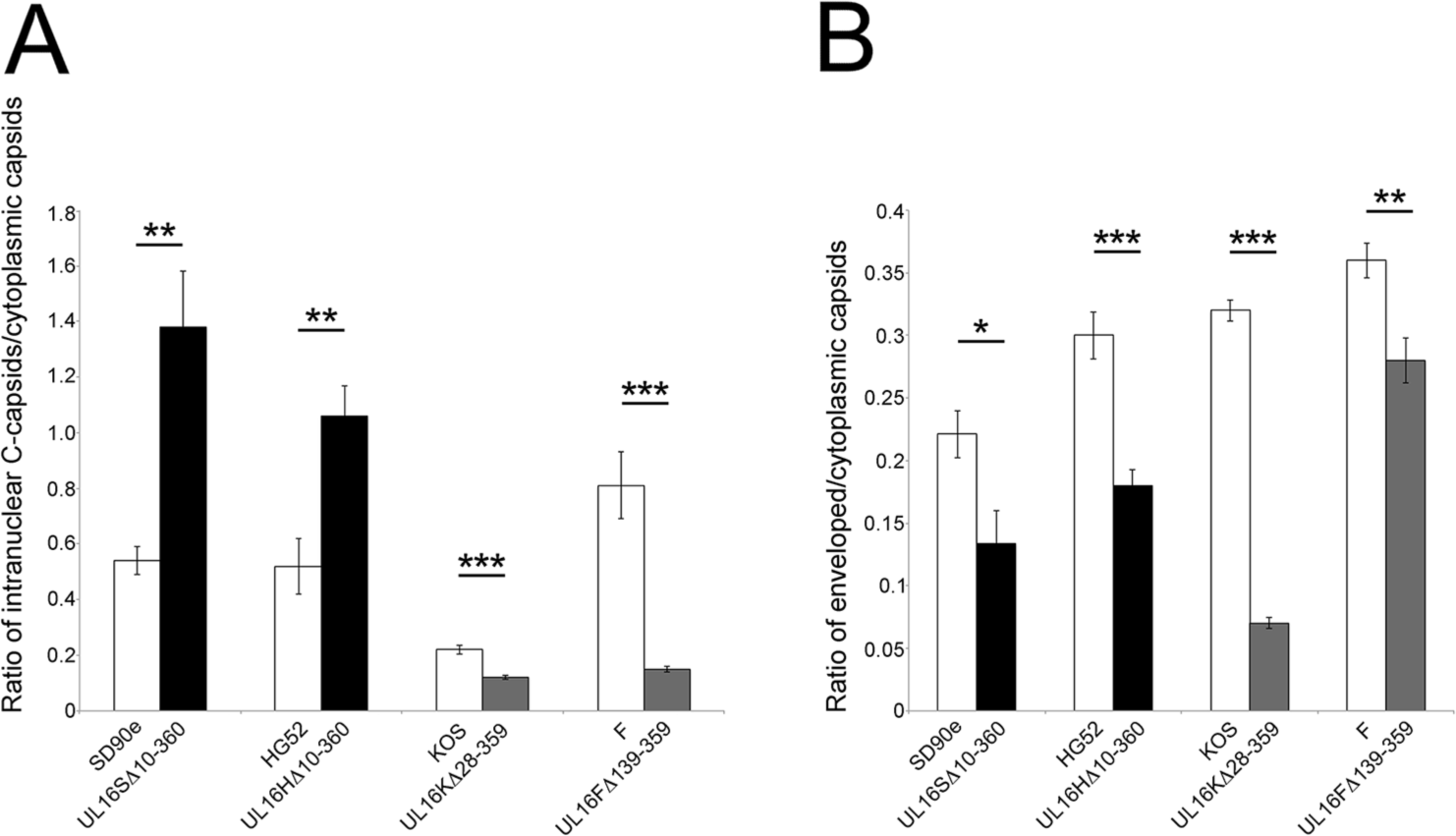
Analysis of capsid distribution in cells infected with HSV *UL16* deletion mutants. (**A**) Ratio of intranuclear C-capsids:cytoplasmic capsids of parental HSV strains and their corresponding *UL16* deletion mutants and were determined. Values were calculated from 10 independent images per strain. Error bars represent standard error of the means. (**B**) Ratio of enveloped:cytoplasmic capsids of parental HSV strains and their corresponding *UL16* deletion mutants were calculated using the same methodology as in (**A**). * * * *P*<0.0001, * * *P*<0.001, * *P*<0.05.

**Table 2.** Quantification of intracellular capsids in Vero cells infected with *UL16* mutant and parental HSV strains^*a*^

| Strain | Total Number of Capsids | Intranuclear Capsids |  | Cytoplasmic Capsids |  | PNS<sup>b</sup> Capsids |
| --- | --- | --- | --- | --- | --- | --- |
|  |  | A+B | C | Non-Enveloped | Enveloped |  |
| SD90e | 948 | 377(39.8<sup>c</sup>) | 96(10.1) | 250(26.4) | 53(5.6) | 172(18.1) |
| UL16SΔ10-360 | 992 | 677(68.2) | 75(7.6) | 159(16) | 13(1.3) | 68(6.9) |
| HG52 | 1429 | 676(47.3) | 107(7.5) | 441(30.9) | 163(11.4) | 42(2.9) |
| UL16HΔ10-360 | 990 | 702(70.9) | 91(9.2) | 156(15.7) | 29(2.9) | 12(1.2) |
| KOS | 850 | 249(29.3) | 75(8.8) | 296(34.8) | 171(20.1) | 59(6.9) |
| UL16KΔ28-359 | 1058 | 506(47.8) | 58(5.5) | 452(42.8) | 35(3.3) | 7(0.7) |
| F | 663 | 217(32.7) | 77(11.6) | 175(26.4) | 126(19) | 68(10.3) |
| UL16FΔ139-359 | 1778 | 892(50.2) | 84(4.7) | 671(37.7) | 99(5.6) | 32(1.8) |

<sup>a</sup> Capsids were counted in different cellular compartments from 10 images/strain derived from multiple sections in two independent experiments.
<sup>b</sup> Perinuclear space (PNS).
<sup>c</sup> Numbers in parentheses represent the percentage of capsids in each category.

Because HSV-1 did not appear to require UL16 for nuclear egress we were interested in determining if the HSV-1 UL16 protein had the capacity to promote the nuclear egress of an HSV-2 *UL16* mutant. To test this, Vero16 and Vero16K cells were infected with UL16SΔ10-360 and UL16KΔ28-359, processed for TEM (Fig. 8) and the TEM data quantified (Fig. 9). Both Vero16 and Vero16K cells were able to support the nuclear egress of UL16SΔ10-360 as evidenced by the appearance of numerous cytoplasmic capsids (Fig.8A, B). Quantification of these data indicated that HSV-2 and HSV-1 UL16 were indistinguishable in their ability to complement the UL16SΔ10-360 nuclear egress defect (Figure 9A). Expression of HSV-1 UL16, but not HSV-2 UL16, modestly, but significantly, promoted the nuclear egress of the UL16KΔ28-359 strain (Figure 9A). As expected, both HSV-2 and HSV-1 UL16 proteins were able to complement secondary envelopment of both HSV species (Fig. 9B). Collectively, these data suggest that HSV-1 encodes a function, missing in HSV-2, that can compensate for nuclear egress in the absence of UL16.

**Fig 8.**
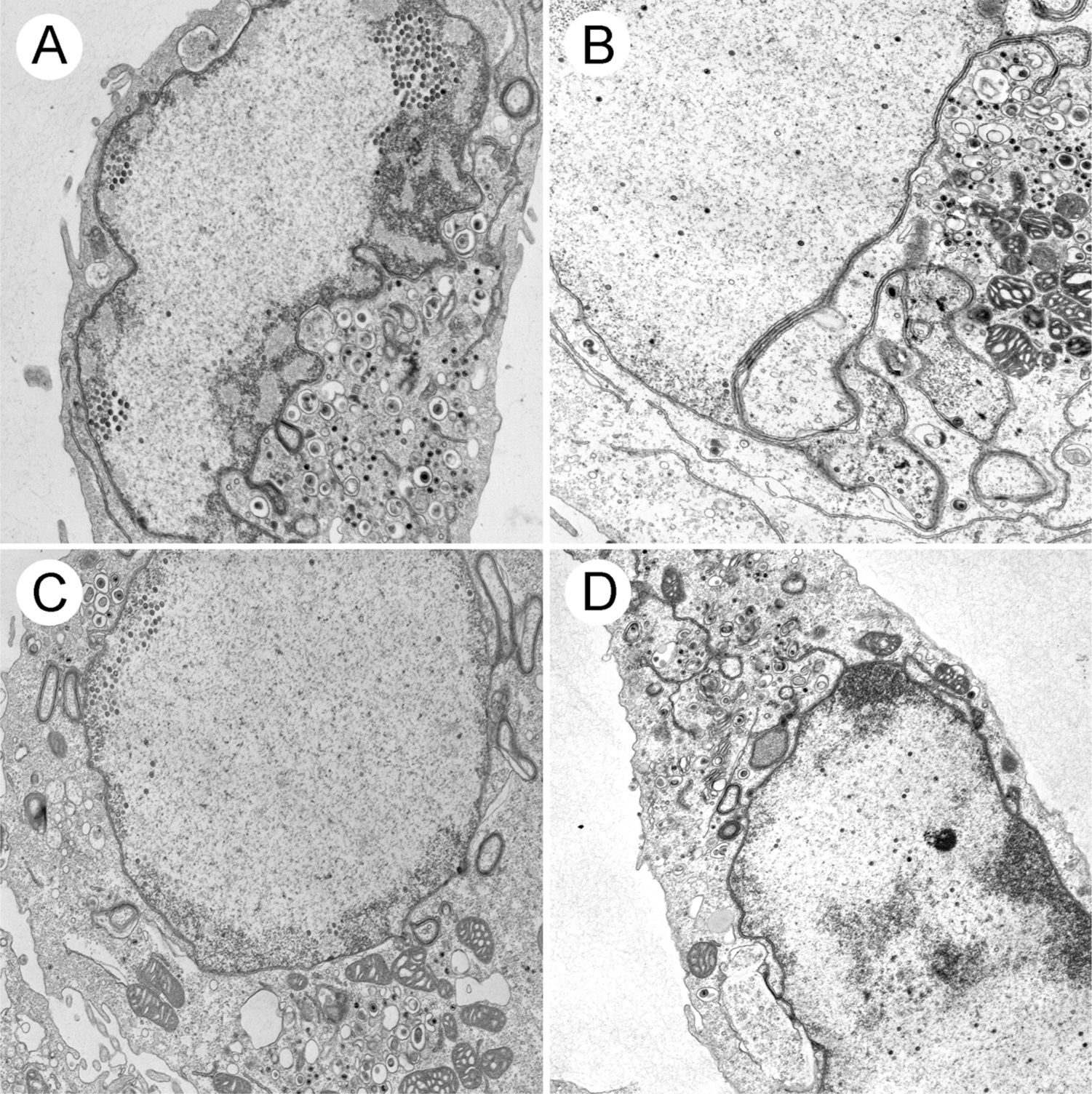
Ultrastructural analysis of trans-complemented *UL16* mutants. Vero16 cells, expressing HSV-2 UL16, and Vero16K cells, expressing HSV-1 UL16, were infected with UL16SΔ10-360 (**A** and **B**) and UL16KΔ28-359 (**C** and **D**), at an MOI of 3. At 16 hpi, cells were fixed and processed for TEM as described in Materials and Methods. Numerous nuclear and cytoplasmic capsids can be observed in all infected cells.

**Fig 9.**
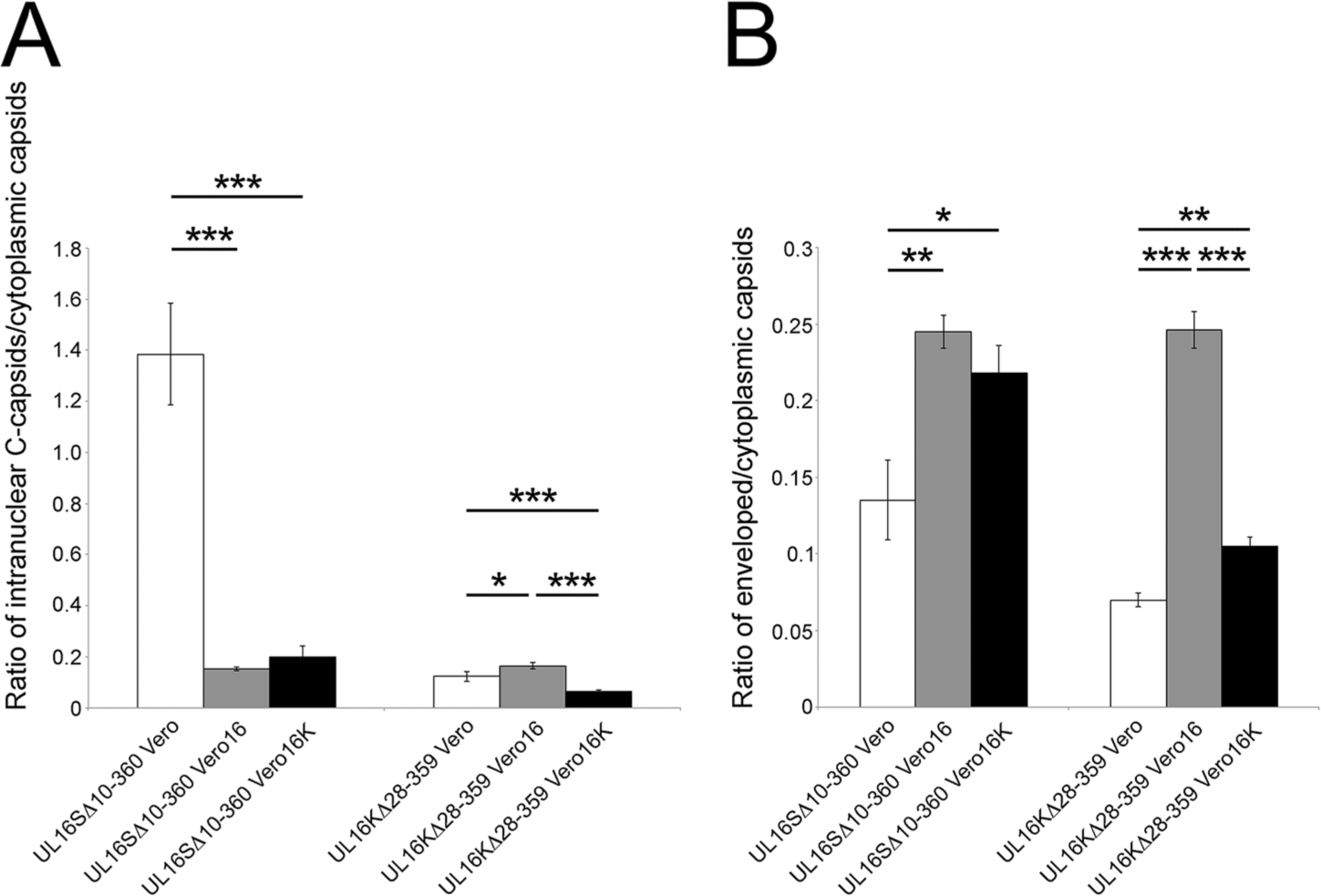
Quantitative analysis of UL16 trans-complementation. (**A**) Ratio of intranuclear C-capsids:cytoplasmic capsids of representative HSV-2 and HSV-1 *UL16* mutants complemented either by HSV-2 (Vero16) or HSV-1 (Vero16K) UL16 protein. Values were calculated from 10 independent images per strain. Error bars represent standard error of the means. (**B**) Ratio of enveloped:cytoplasmic capsids of representative HSV-2 and HSV-1 *UL16* mutants complemented either by HSV-2 or HSV-1 UL16 protein were calculated using the same methodology as in (**A**). *** *P*<0.0001, ** *P*<0.001, * *P*<0.05.

## Discussion

Here we describe the analysis of *UL16* deletion mutants derived from four HSV strains. The strategy used to construct these strains utilized CRISPR/Cas9 mutagenesis which was both efficient and rapid. Our approach utilized two gRNAs towards the *UL16* locus simultaneously. Cleavage of the *UL16* gene at the sites directed by the gRNAs and subsequent repair of the lesion by non-homologous end joining (NHEJ) resulted in the isolation of a variety of mutants; some having in-frame deletions and others with frame-shifts after the 5’ gRNA directed cleavage. For the analysis presented here we chose to select a variety of *UL16* mutants for further study. It is noteworthy that all HSV-2 *UL16* mutants isolated displayed a similar phenotype. In the case of HSV-1, however, one of the F *UL16* mutants, UL16FFS27/3, was an outlier insofar as its replication was reduced much more severely than other HSV-1 *UL16* mutants (Fig. 3D). It is not clear why UL16FFS27/3 grows as poorly as it does, however, it is noteworthy that it forms smaller plaques on complementing cells than the other HSV-1 *UL16* mutants (Fig.4B) raising the possibility that additional mutations outside the UL16 locus were introduced during its isolation. Alternatively, the N-terminal fragment of UL16, predicted to be produced by UL16FFS27/3, might act as a dominant negative protein resulting in the inhibition of both cell-to-cell spread and virus replication. Clearly, more work is required to determine the cause of the UL16FFS27/3 cell-to-cell spread and replication phenotypes. Because of these caveats we eliminated this strain from subsequent ultrastructural analyses.

Our kinetic analysis of *UL16* mutant replication revealed that HSV-2 *UL16* mutants had roughly 50 to 100-fold reductions in virus replication while HSV-1 *UL16* mutants, with the exception of UL16FFS27/3 (see above), had approximately 10-fold reductions (Fig. 3). These results are consistent with previous findings suggesting HSV-2 and HSV-1 have differential requirements for UL16 (6-8). Despite replicating better than HSV-2 *UL16* deletion mutants (Fig. 3), HSV-1 *UL16* mutants consistently formed slightly smaller plaques relative to their parental strains than the HSV-2 *UL16* mutants (Fig. 2). These findings may suggest that HSV-1 has a greater reliance on UL16 for cell-to-cell spread of infection than does HSV-2. Along these lines, Wills and colleagues have documented an interaction between the N-terminus of HSV-1 UL16 and the cytoplasmic tail of glycoprotein gE (14) and that a complex formed by UL16, UL11 and UL21 on the gE cytoplasmic tail is important for normal glycosylation of gE, trafficking of gE to the cell surface and cell-to-cell spread of infection (14, 15). The existence of such interactions and their potential roles in the spread of HSV-2 infection have yet to be determined. Perhaps such interactions are not required for efficient cell-to-cell spread in HSV-2 infected cells and therefore might explain the differences in relative plaque sizes observed. In support of this idea, it is noteworthy that the N-terminus of UL16 is less conserved between HSV-2 and HSV-1 than the remainder of the protein (16) and our preliminary investigations suggest that gE glycosylation is unperturbed in cells infected with HSV-2 Δ16 (data not shown).

Our trans-complementation plaque assays revealed that HSV-1 UL16 can rescue plaque formation of HSV-2 *UL16* mutants and vice versa (Fig. 4). Furthermore, TEM analysis revealed that HSV-1 UL16 can promote the nuclear egress of HSV-2, despite not being required for HSV-1 nuclear egress (Fig. 8 and 9A). Our findings also indicate that both HSV-2 and HSV-1 UL16 function in secondary envelopment and that these proteins are trans-complementary for this process (Fig.8 and 9B). Perhaps there are similarities in the processes of primary and secondary envelopment and HSV-1 UL16 is able to function in primary envelopment in the context of HSV-2 infection. The observation that HSV-1 and HSV-2 UL16 molecules can complement each other suggests that the genetic basis for the species-specific activities of UL16 lie outside the *UL16* locus.

Importantly, these findings do not fully explain the reductions in virus replication observed for all *UL16* mutant strains. The explanation for the magnitude of the replication deficiencies observed for *UL16* mutants are certainly multifactorial. The functions of HSV-1 UL16 in cell-to-cell spread of infection have been well documented (14, 15). Additionally, previous studies on HSV-2 UL16 have implied a role for UL16 in viral DNA packaging into capsids (16). Moreover, the proportion of perinuclear virions compared to parental strains was reduced in all *UL16* mutants examined (Table 2). Taken together, these findings suggest that UL16 influences multiple stages of virion morphogenesis.

The goal of this study was to resolve an apparent discrepancy between the functions of HSV-2 and HSV-1 UL16 during virus maturation. We have conclusively demonstrated that UL16 is important for HSV-2 nuclear egress in multiple strains (186, SD90e and HG52). Additionally, it is clear that multiple strains of HSV-2 and HSV-1 rely on UL16 for efficient secondary envelopment. Despite important differences in primary and secondary envelopment, such as the well-characterized functions of the nuclear egress complex in primary envelopment, these findings raise the intriguing possibility that some aspects of primary and secondary envelopment may be more similar than previously appreciated.

## Materials and Methods

### Viruses and cells

HSV-2 strains 186 and SD90e were kind gifts from David Knipe, Harvard University. The construction of HSV-2 186 strain UL16 knockout (Δ16) was described previously (6). HSV-2 strain HG52 was kindly provided by Aidan Dolan and Duncan McGeoch, University of Glasgow. HSV-1 strains F and KOS were generously provided by Lynn Enquist, Princeton University. African green monkey kidney cells (Vero) and human embryonic kidney 293T cells were acquired from the ATCC. Phoenix-AMPHO cells were generously provided by Craig McCormick, Dalhousie University. The murine L fibroblast cell line was a kind gift from Frank Tufaro, University of British Columbia. All cell lines were cultured in Dulbecco’s modified Eagle’s medium (DMEM) supplemented with 10% fetal bovine serum (FBS), 1% penicillin-streptomycin, and 1% GlutaMAX and grown at 37°C in a 5% CO_2_ environment.

UL16 expressing cell lines were isolated by retroviral transduction using an amphotropic Phoenix-Moloney murine leukemia virus system described previously (17). In brief, plasmids pBMN-IP-UL16 or pBMN-IP-UL16K (see below) were transfected into Phoenix-AMPHO cells to produce the retroviruses. HSV-2 UL16 expressing cell lines (Vero16 and 293T16) and HSV-1 UL16 expressing cell lines (L16K and Vero16K) were isolated by transducing either Vero, 293T or L cells with the corresponding amphotrophic retroviruses, and were selected using 2 µg/mL puromycin (InvivoGen) 48 hrs after transduction. To confirm UL16 expression, cell extracts were prepared and analyzed by Western blotting using HSV-2 or HSV-1 UL16 antiserum (Sup Fig. 1).

### Antibodies

Chicken polyclonal antiserum against HSV-2 UL16 (6) was used for Western blotting at a dilution of 1: 200, mouse monoclonal antibody against HSV-2 ICP27 (Virusys) was used for Western blotting at a dilution of 1: 1,000. Rabbit polyclonal antiserum against HSV-1 UL16 was a kind gift from John Wills, The Pennsylvania State University College of Medicine (18), which was used for Western blotting at a dilution of 1: 3,000. Rat polyclonal antiserum against Us3 (19) was used for indirect immunofluorescence microscopy at a dilution of 1: 1,000 and mouse monoclonal antibody against β-actin (Sigma) was used for Western blotting at a dilution of 1: 2,000. Alexa Fluor 568-conjugated goat anti-rat immunoglobulin G monoclonal antibody (Invitrogen Molecular Probes) was used at a dilution of 1:500 for immunofluorescence microscopy. Horseradish peroxidase-conjugated goat antimouse IgG, horseradish peroxidase-conjugated goat anti-chicken IgY, horseradish peroxidase-conjugated rabbit anti-rat IgG and horseradish peroxidase-conjugated goat antirabbit IgG (Sigma) were used for Western blotting at dilutions of 1: 10,000, 1: 30,000, 1: 80,000 and 1: 5,000, respectively.

### Plasmid construction

pBMN-IP-UL16 encoding HSV-2 UL16 was constructed previously (6). To construct pBMN-IP-UL16K, UL16 KOS sequences were amplified from HSV-1 KOS genomic DNA by PCR using the forward primer 5’-GACT*GAATTC*ATGG CGCAGCTGGGAC-3’ containing an EcoRI restriction site (italics) and reverse primer 5’-GACT*CTCGAG*TTATTCGGGATCGCTTG-3’ containing a XhoI restriction site (italics). The PCR product was digested with EcoRI and XhoI and ligated into similarly digested pBMN-IP (a kind gift of Craig McCormick, Dalhousie University) to yield pBMN-IP-UL16K.

Guide RNAs (gRNAs) used for producing the *UL16* mutant strains were expressed from the guide RNA-Cas9 expression plasmid pX330-U6-Chimeric_BB-CBh-hSpCas9, a gift from Feng Zhang (Addgene plasmid 42230) (20). To construct these gRNA expression plasmids the top-strand oligonucleotide was annealed to the bottom-strand oligonucleotide (Table 1) and the double-stranded product was cloned into pX330-U6-Chimeric_BB-CBh-hSpCas9, that had been digested with BbsI. Three different *UL16* gRNAs were designed, for both HSV-1 and HSV-2, to produce different sized deletions within *UL16*.

**Table 1.**
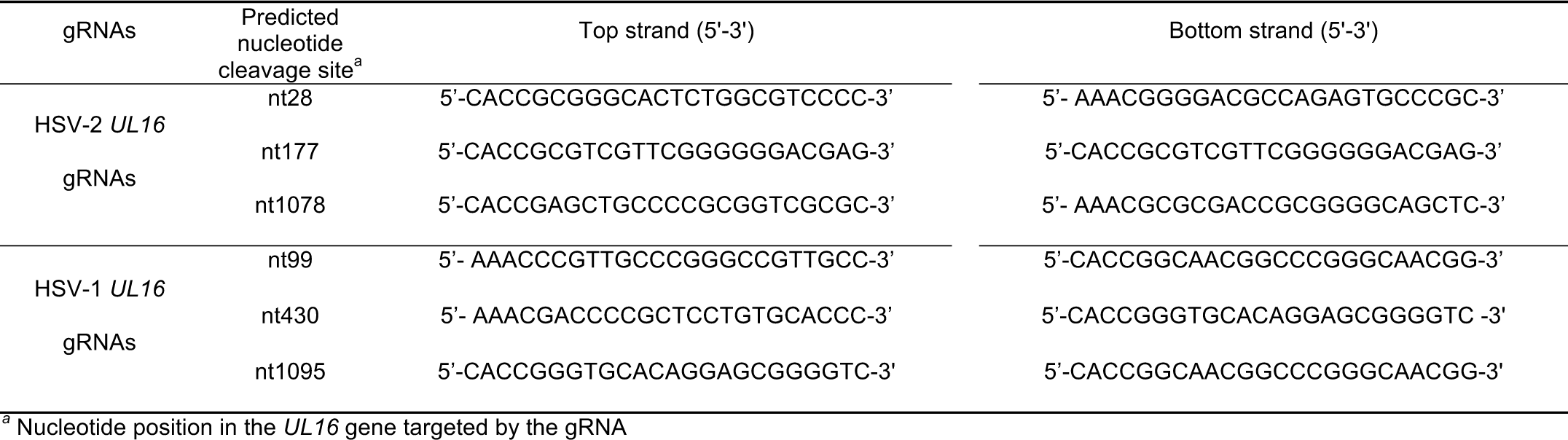
Oligonucleotides used to produce HSV-2 and HSV-1 *UL16* gRNAs

### CRISPR/Cas9 mutagenesis of the *UL16* locus

A similar approach to that used by Xu and colleagues for the construction of PRV mutants was utilized (21). Viral DNA of each strain (SD90e, HG52, KOS, or F) was purified as described previously (22). 293T16 cells growing in 100-mm dishes were co-transfected with 16μg of purified viral genomic DNA along with 1µg each of two *UL16* guide RNA expression plasmids using a calcium phosphate co-precipitation method (23). 24h after transfection, the culture medium was replaced with semisolid medium containing 0.5% methyl cellulose to allow for plaque formation. Five to six days later, plaques were picked. Viral DNA isolated from a portion of the picked plaque was used for screening for *UL16* deletions by PCR. The UL16 locus from viruses bearing *UL16* deletions were sequenced in their entirety to determine the precise nature of the *UL16* mutations introduced. Roughly 50% of plaques picked had *UL16* deletions or frame shift mutations.

### Plaque size determination

Monolayers of Vero cells were prepared on 35-mm glass bottom dishes (MatTek), and infected with virus at a multiplicity of infection (MOI) of 0.005. Plaques were allowed to form for 24h prior to fixation and processing for indirect immunofluorescence microscopy (6) using antisera against the HSV Us3 protein (19). Images of plaques were captured on a Nikon TE200 inverted epifluorescence microscope using a 10X objective and a cooled CCD camera. To quantify these results, the numbers of pixels in the area of each plaque were counted using Image-Pro 6.3 software. Results shown were derived from 40 distinct plaques per strain.

### Transmission electron microscopy (TEM)

Vero cells growing in 100-mm dishes were infected with virus at an MOI of 3 and processed for TEM at 16 hpi. Infected cells were rinsed with PBS three times before fixing in 1.5 mL of 2.5% glutaraldehyde in 0.1 M sodium cacodylate buffer (pH 7.4) for 60 mins. Cell were collected by scraping into fixative and centrifugation at 300×g for 5 min. Cell pellets were carefully enrobed in an equal volume of molten 5% low-melting point agarose and allowed to cool. Specimens embedded in agarose were incubated in 2.5% glutaraldehyde in 0.1 M sodium cacodylate buffer (pH 7.4) for 1.5 hrs and post-fixed in 1% osmium tetroxide for 1 hr. The fixed cells in agarose were rinsed with distilled water 3 times and stained in 0.5% uranyl acetate overnight before dehydration in ascending grades of ethanol (30%-100%). Samples were transitioned from ethanol to infiltration with propylene oxide and embedded in Embed-812 hard resin (Electron Microscopy Sciences). Blocks were sectioned at 50-60 nm and stained with uranyl acetate and Reynolds’ lead citrate. Images were collected using a Hitachi H-7000 transmission electron microscope operating at 75kV.

## Acknowledgements

This work was supported by the Canadian Institutes of Health Research operating grant 93804, Natural Sciences and Engineering Council of Canada Discovery Grant 418719 and Canada Foundation for Innovation award 16389 to BWB. JG was supported in part by an award from the China Scholarship Council. We are grateful to Kelsey Wan for help with viral DNA preparation, Laura Ruhge and Greg Smith for advice with sample preparation for electron microscopy and Renée Finnen for critical reading of the manuscript.

